# Protein-Protein Interaction Network Analysis and Identification of Key Players in nor-NOHA and NOHA Mediated Pathways for Treatment of Cancer through Arginase Inhibition: Insights from Systems Biology

**DOI:** 10.1101/281030

**Authors:** Ishtiaque Ahammad

## Abstract

L-arginine is involved in a number of biological processes in our bodies. Metabolism of L-arginine by the enzyme arginase has been found to be associated with cancer cell proliferation. Arginase inhibition has been proposed as a potential therapeutic means to inhibit this process. N-hydroxy-nor-L-Arg (nor-NOHA) and N (omega)-hydroxy-L-arginine (NOHA) has shown promise in inhibiting cancer progression through arginase inhibition. In this study, nor-NOHA and NOHA-associated genes and proteins were analyzed with several Bioinformatics and Systems Biology tools to identify the associated pathways and the key players involved so that a more comprehensive view of the molecular mechanisms including the regulatory mechanisms can be achieved and more potential targets for treatment of cancer can be discovered. Based on the analyses carried out, 3 significant modules have been identified from the PPI network. Five pathways/processes have been found to be significantly associated with nor-NOHA and NOHA associated genes. Out of the 1996 proteins in the PPI network, 4 have been identified as hub proteins-SOD, SOD1, AMD1, and NOS2. These 4 proteins have been implicated in cancer by other studies. Thus, this study provided further validation into the claim of these 4 proteins being potential targets for cancer treatment.

## Introduction

L-arginine, a basic amino acid, has a major role in a number of systems in our bodies including the immune system, particularly on the proliferation of T lymphocytes [1]. The enzyme arginase metabolizes L-arginine to L-ornithine which is important in the biochemical pathways involved in cell proliferation [2]. Two isoforms of arginase exists: arginase I, and arginase II. Arginase I is a cytosolic enzyme which is primarily found in hepatocytes, erythrocytes, and granulocytes [3], and Arginase II is found in the mitochondria of various tissues, including kidney, brain, and prostate [5], [6]. Arginase II activity has been shown to be increased in breast, colon, and prostate cancer [7], [8]

Arginase inhibition has been found to suppress proliferation of breast cancer cells. Therefore it is being considered as a new therapeutic target for suppressing breast cancer cell growth. In tumor cells with elevated arginase activity, arginase inhibitors can be a potential therapeutic option for treating cancer [9].

N(omega)-hydroxy-L-arginine (NOHA) has been found to selectively inhibit cell proliferation and induce apoptosis in high arginase expressing MDA-MB-468 cells [7]. NOHA is also involved in the modulation of T cell receptor CD3zeta which undergoes down-regulation in cancer [7].

Injection of N-hydroxy-nor-L-Arg (nor-NOHA) blocked growth of lung cancer cells in mice [10]. In another study, nor-NOHA significantly (P = 0.01) reduced arginase II activity and suppressed growth of cells with high arginase activity in renal carcinoma [9]

The purpose of this study was to identify crucial genes and proteins involved in the nor-NOHA and NOHA mediated pathways in order to shed more light on their mechanism of action and the key players involved. For this, nor-NOHA and NOHA-associated genes from several databases was identified. And multiple Bioinformatics and Systems Biology tools have been used to identify more nor-NOHA and NOHA-associated key genes. They included pathway enrichment analysis, protein-protein interaction (PPI) network, module analysis and transcriptional regulatory network analysis.

## Methods

### Nor-NOHA and NOHA-Associated Genes

“nor-NOHA” and “NOHA” were used as keywords to search in three databases for genes associated with these compounds and the results were combined to build a set of Nor-NOHA and NOHA-associated genes. The three databases used were-

- GeneCards (version 3.0), a searchable, integrative database which provides comprehensive information on all annotated and predicted human genes [11].
- Search Tool for Interacting Chemicals (STICH, version 5.0), a database of known and predicted interactions between chemicals and proteins [12]
- Comparative Toxicogenomics Database (CTD), which provides manually curated information concerning chemical–gene/protein interactions, chemical–disease as well as gene–disease relationships was used [13].

### PPI Network

Two databases were combined to predict the Protein-Protein Interaction (PPI) pairs among the nor-NOHA and NOHA-associated genes. The databases used were-

- Biological General Repository for Interaction Datasets (BioGRID, version 3.4, https://wiki.thebiogrid.org/) [14]
- The Molecular Interaction Database (MINT, 2012 update, https://mint.bio.uniroma2.it/) [15]

Using the software Cytoscape (http://www.cytoscape.org) [16], a PPI network was visualized for Nor-NOHA and NOHA-associated genes.

Using the CytoNCA plug-in [17] (version 2.1.6, http://apps.cytoscape.org/apps/cytonca) in Cytoscape, degree centrality (DC), betweenness centrality (BC), and closeness centrality of the nodes of the PPI network were subjected to analysis to identify the hub proteins [18]. “Without weight.” was set as the parameter.

### Module Analysis

For module analysis of the PPI network, MCODE plug-in [19] (version 1.4.2; http://apps.cytoscape.org/apps/mcode; parameters set as degree cut-off = 2, maximum depth = 100, node score cut-off = 0.2, and -core = 2) in Cytoscape was used.

### Pathway Enrichment Analysis

KEGG pathway enrichment analysis for the nodes of top modules was carried out with JEPETTO plug-in [20] in Cytoscape.

### Transcriptional Regulatory Network Construction

Transcription factors (TFs) among nor-NOHA and NOHA-associated genes were searched and then their targets were identified using the transcriptional regulatory relationships unravelled by a sentence-based text-mining (TRRUST, http://www.grnpedia.org/trrust/) [21] database.

Finally, a transcriptional regulatory network of the hub proteins was constructed using Cytoscape [16].

### Results and Discussion

In this study, a total of 19 nor-NOHA and NOHA-associated genes were identified from GeneCards, STICH and CTD databases (Table 1) and their protein-protein interaction network was constructed (Figure 1). Based on Betweenness Centrality, Closeness Centrality, and Degree Centrality scores SOD, SOD1, AMD1, and NOS2 were established as hub nodes in the Protein-Protein Interaction network of these genes and their interactions (Table 2). Three distinct modules (modules 1, 2 and 3) of the PPI network were identified (Figure 2). Arginine and proline metabolism, tryptophan metabolism, nucleotide excision repair, amyotrophic lateral sclerosis (ALS) and peroxisome were the top 5 pathways/processes the proteins in these modules were found to be significantly involved in (Table 3). Only 2 out of the 4 hub proteins, namely SOD1 and NOS2 were present in the TRRUST database of transcription factors (Table 5). So a transcriptional regulatory network of them was constructed (Figure 3). SOD1 has been found to be the target of transcription factors SP1, KLF4, EGR1, PPARD and NFE2L2 while NOS2 was found to be the target of transcription factors XBP1, JUN, JUND, STAT1, APC, XBP1, and RELA.

**Table 1:** Nor-NOHA and NOHA associated genes

| Nor-NOHA associated genes | NOHA associated genes |
| --- | --- |
| NOS3 | NOS1 |
| ARG2 | MARC2 |
| NOS2 | NOS2 |
| CD247 | MARC1 |
| NOS1 | NOS3 |
| SOD1 | ARG1 |
| DDAH1 | CYCS |
| IL6 | ODC1 |
| AGMAT | SOD1 |
|  | AMD1 |
|  | CAT |
|  | ARG2 |
|  | STX8 |
|  | UROD |
|  | AGMAT |

**Table 2:**
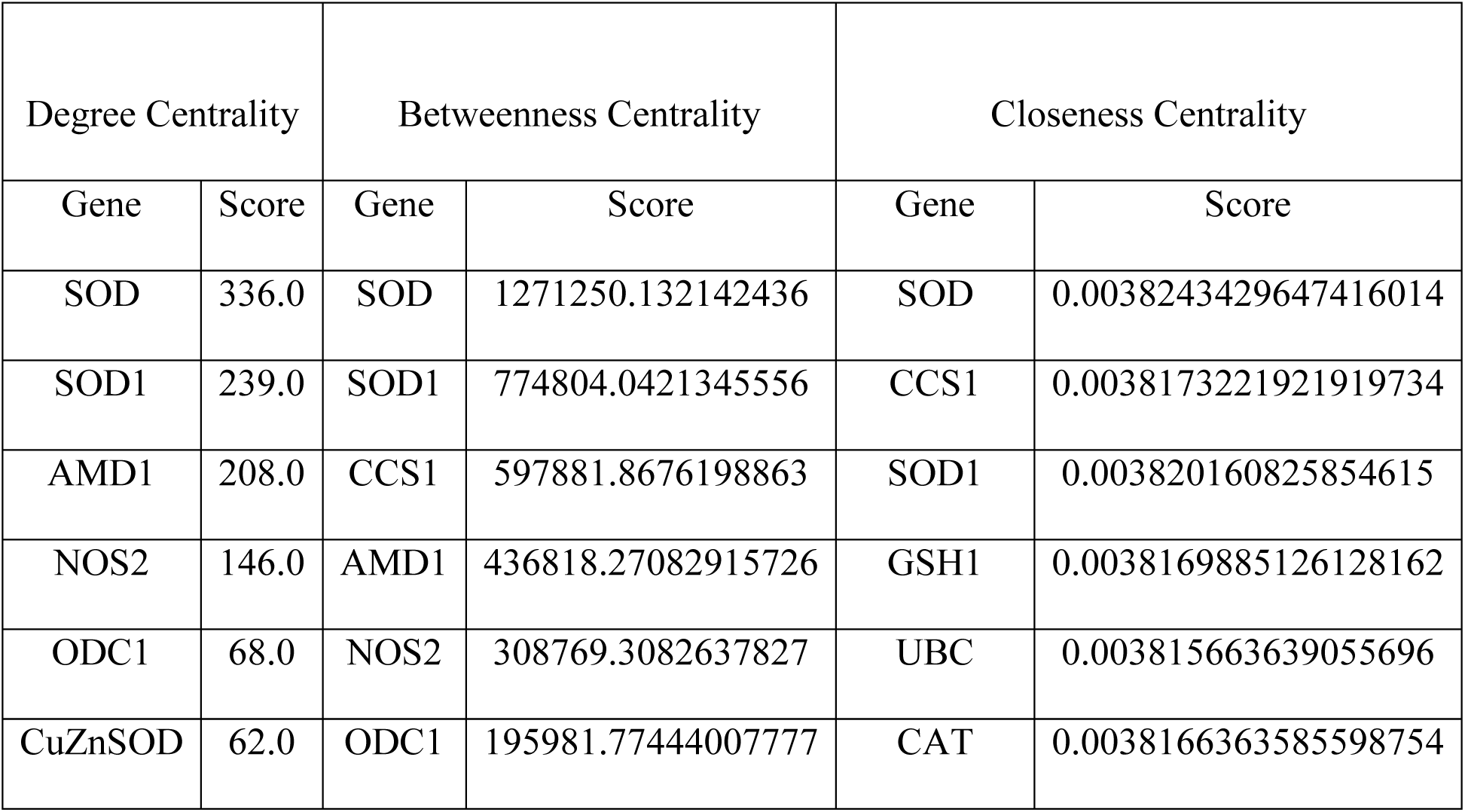
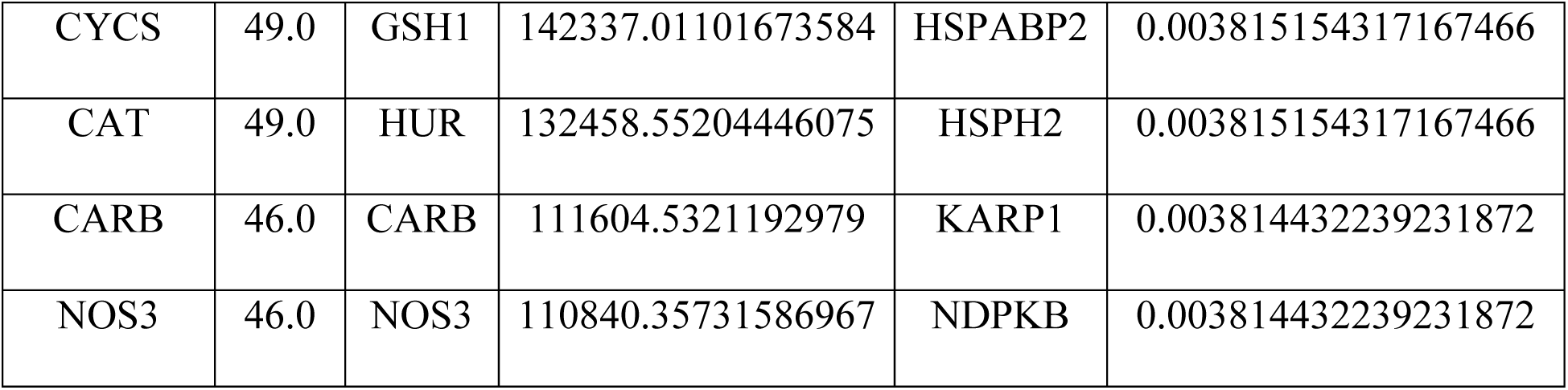
Hub proteins in the PPI network according to Degree Centrality, Betweenness Centrality and Closeness Centrality using CytoNCA

| Degree Centrality |  | Betweenness Centrality |  | Closeness Centrality |  |
| --- | --- | --- | --- | --- | --- |
| Gene | Score | Gene | Score | Gene | Score |
| SOD | 336.0 | SOD | 1271250.132142436 | SOD | 0.0038243429647416014 |
| SOD1 | 239.0 | SOD1 | 774804.0421345556 | CCS1 | 0.0038173221921919734 |
| AMD1 | 208.0 | CCS1 | 597881.8676198863 | SOD1 | 0.003820160825854615 |
| NOS2 | 146.0 | AMD1 | 436818.27082915726 | GSH1 | 0.0038169885126128162 |
| ODC1 | 68.0 | NOS2 | 308769.3082637827 | UBC | 0.003815663639055696 |
| CuZnSOD | 62.0 | ODC1 | 195981.77444007777 | CAT | 0.0038166363585598754 |
| CYCS | 49.0 | GSH1 | 142337.01101673584 | HSPABP2 | 0.003815154317167466 |
| CAT | 49.0 | HUR | 132458.55204446075 | HSPH2 | 0.003815154317167466 |
| CARB | 46.0 | CARB | 111604.5321192979 | KARP1 | 0.003814432239231872 |
| NOS3 | 46.0 | NOS3 | 110840.35731586967 | NDPKB | 0.003814432239231872 |

**Table 3:** KEGG pathway enrichment analysis result for the nodes of top modules

| Pathway or Process | XD-score | q-value | Overlap/Size |
| --- | --- | --- | --- |
| Arginine and proline metabolism | 0.21151 | 0.09869 | 2/37 |
| Tryptophan metabolism | 0.14914 | 1.00000 | 1/26 |
| Nucleotide excision repair | 0.09053 | 1.00000 | 1/42 |
| Amyotrophic lateral sclerosis (ALS) | 0.08040 | 1.00000 | 1/47 |
| Peroxisome | 0.06672 | 1.00000 | 1/56 |
| VEGF signaling pathway | 0.05981 | 1.00000 | 1/62 |
| Amoebiasis | 0.03611 | 1.00000 | 1/98 |
| Protein processing in endoplasmic reticulum | 0.02407 | 1.00000 | 1/139 |
| Calcium signaling pathway | 0.02161 | 1.00000 | 1/152 |
| Metabolic pathways | 0.01404 | 1.00000 | 3/640 |

**Table 4:**
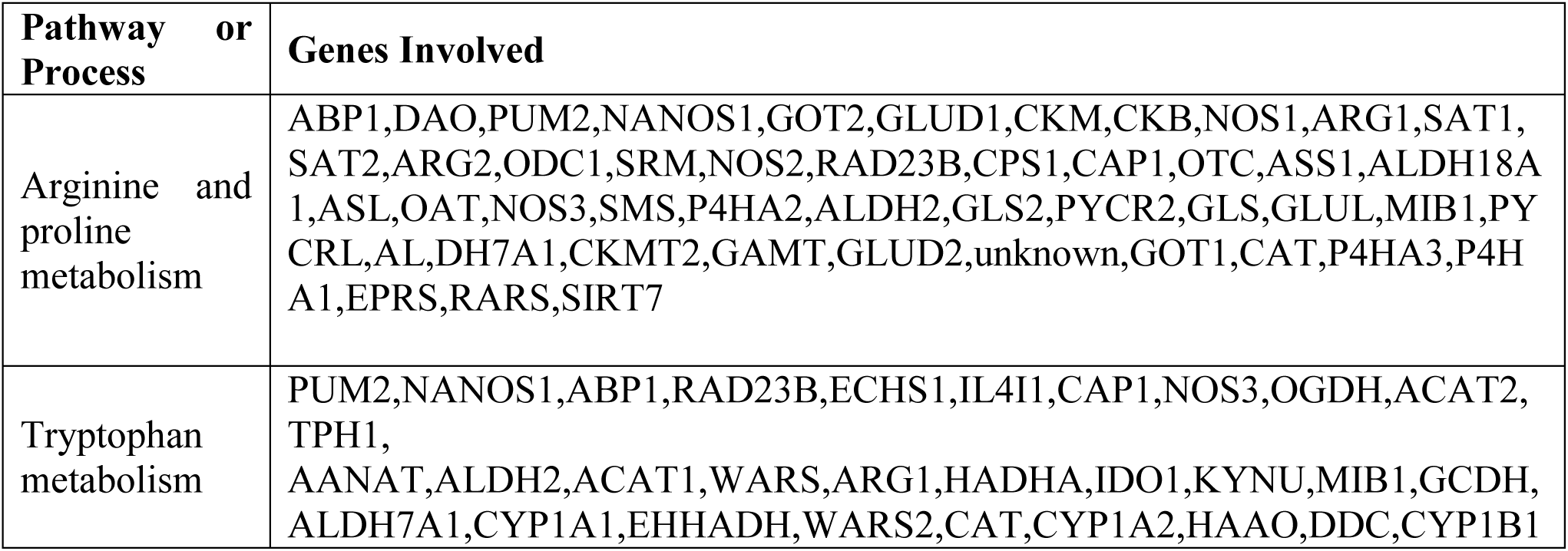
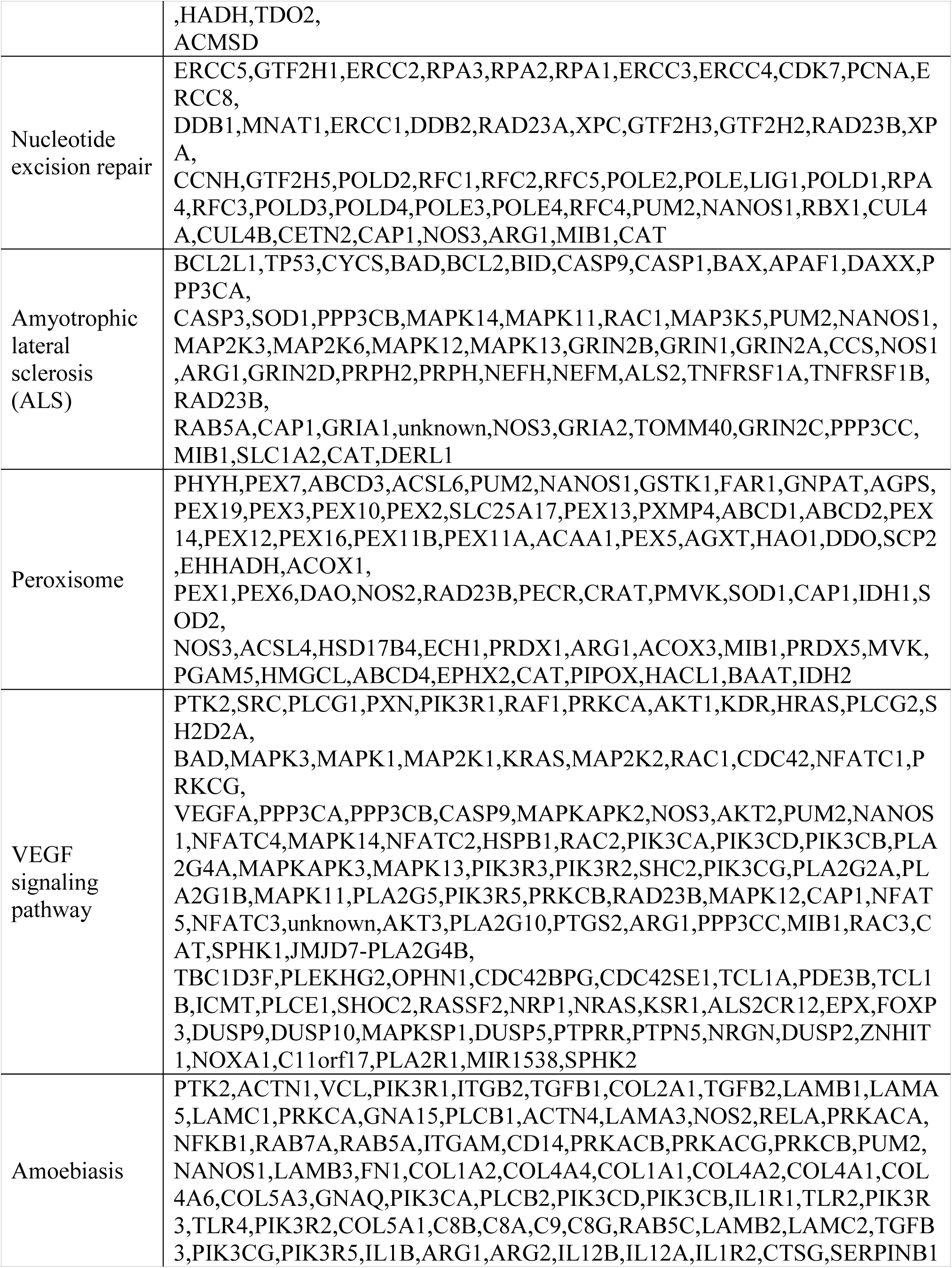
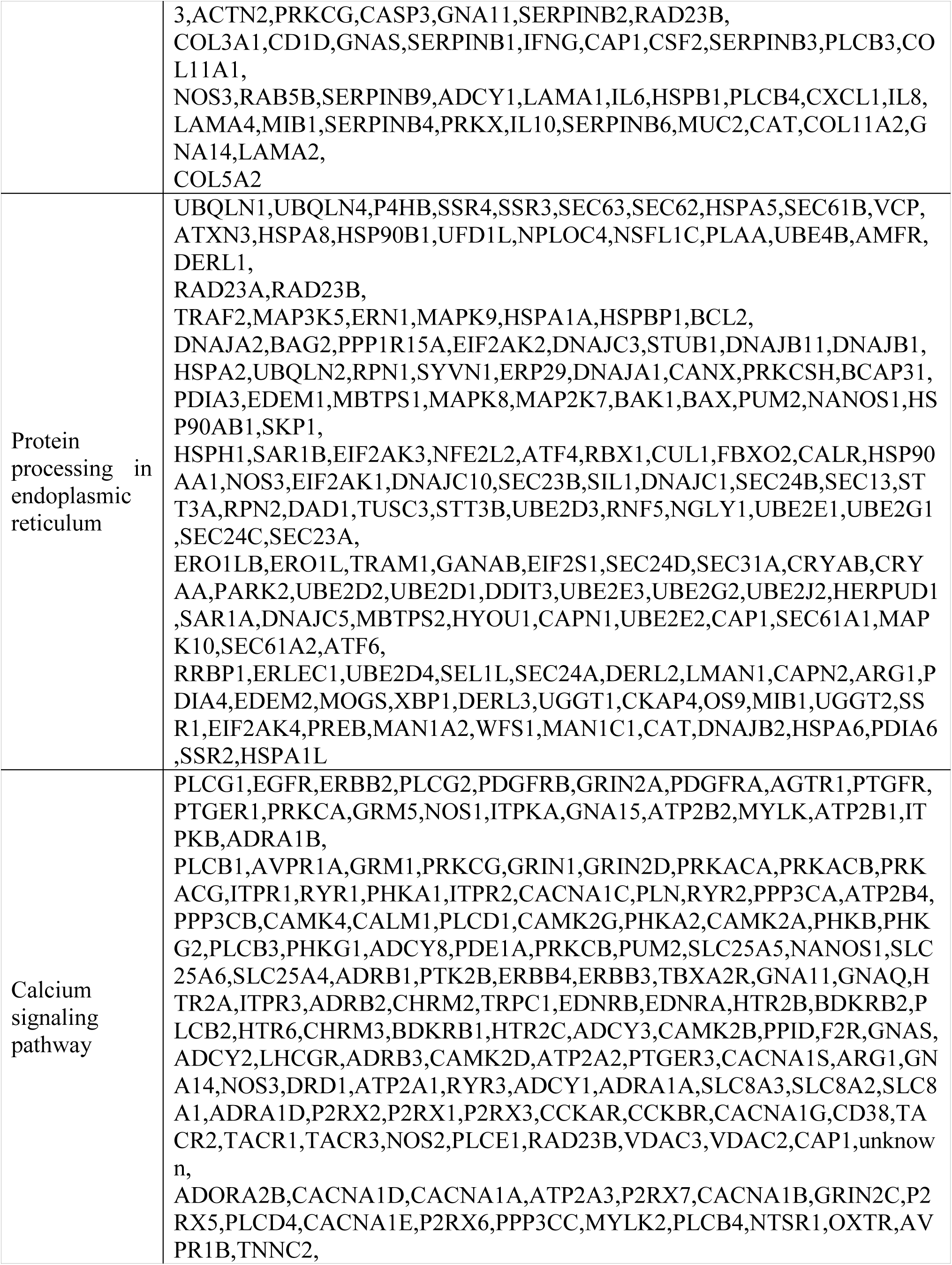
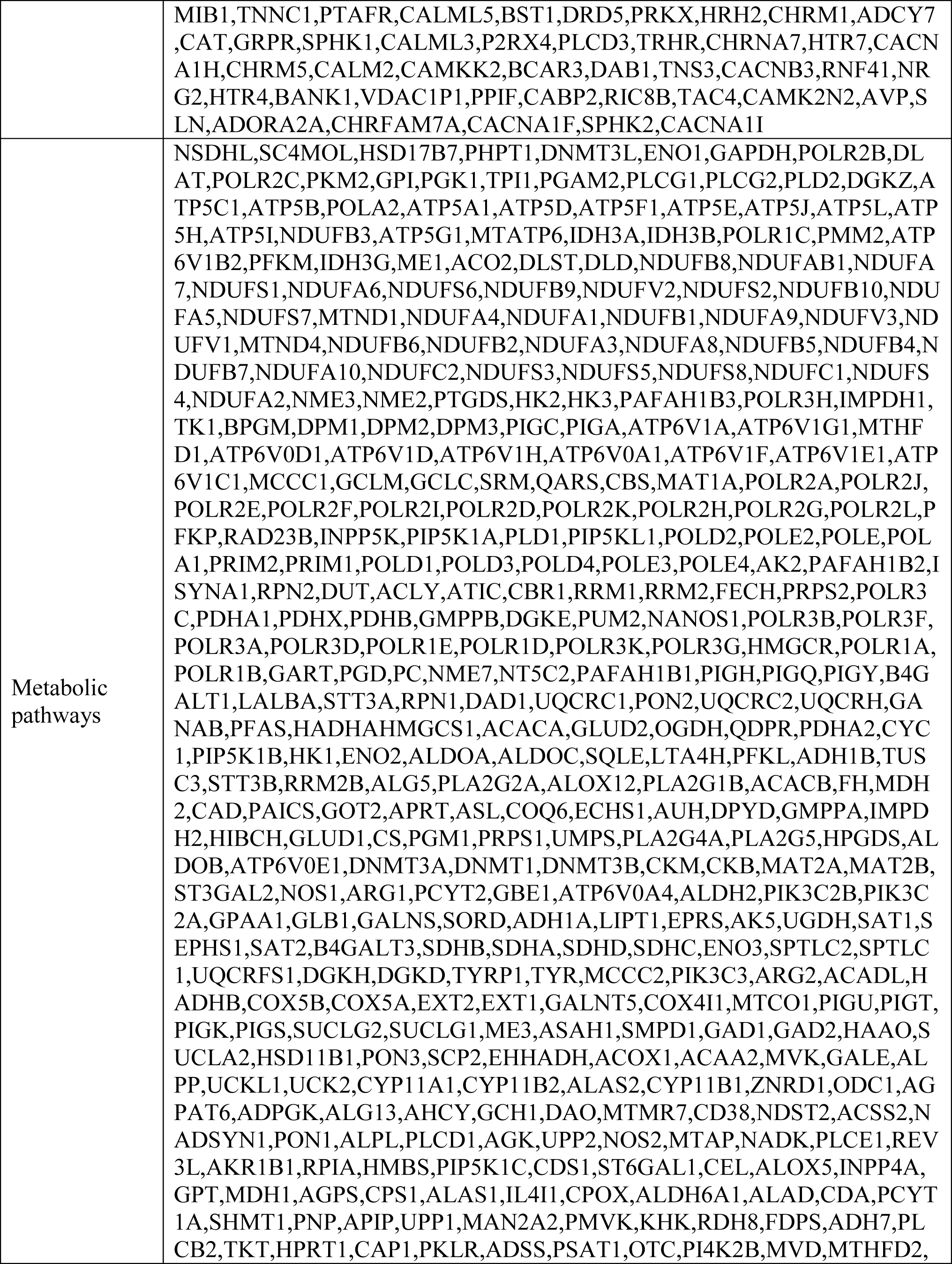
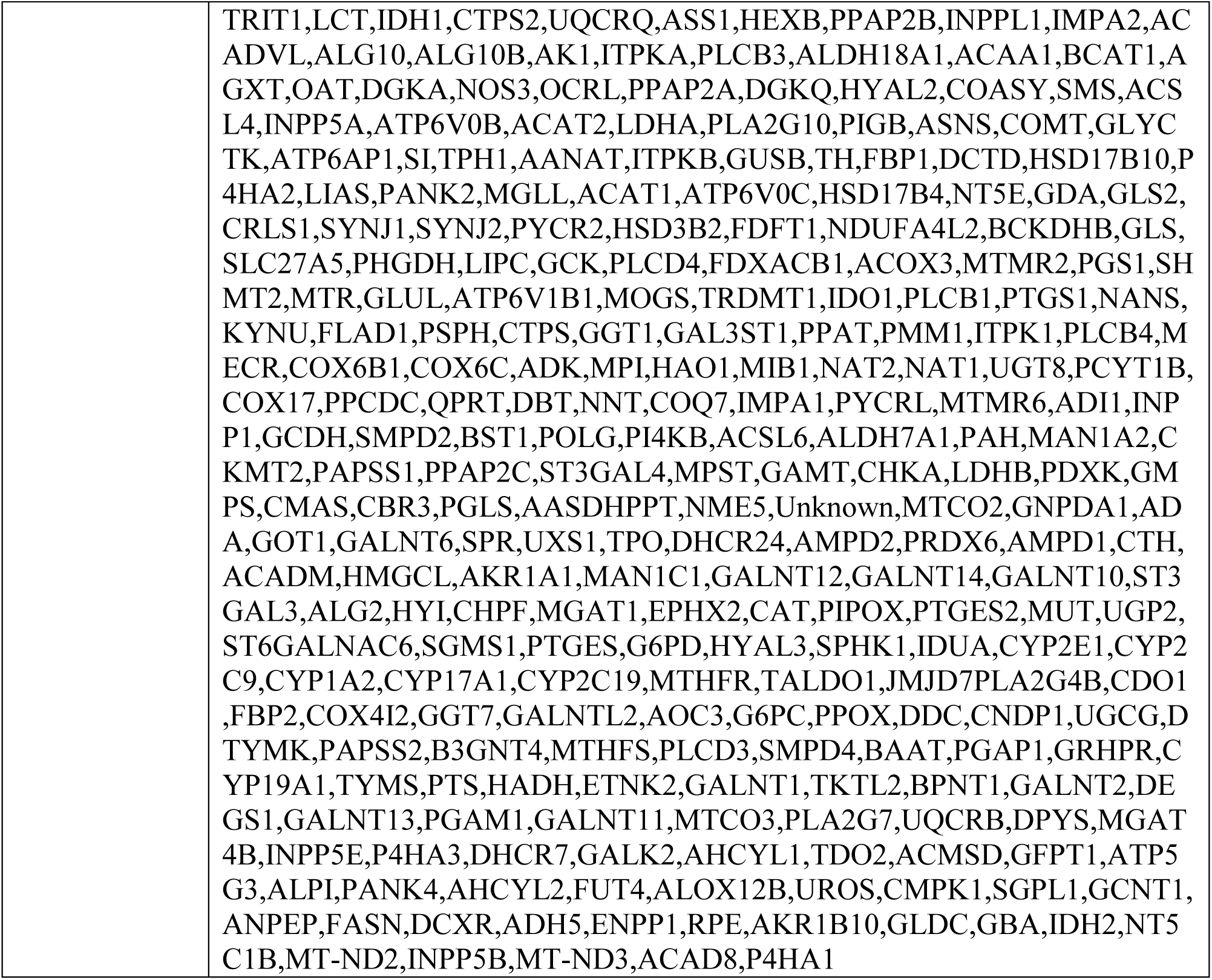
Genes involved in the top 10 enriched KEGG pathways

**Table 5:** Transcription factors targeting nor-NOHA and NOHA associated genes

| # | Key TF | Description | Target genes | Mode of action |
| --- | --- | --- | --- | --- |
| 1 | SP1 | Sp1 transcription factor | CAT | Unknown |
|  |  |  | IL6 | Activation |
|  |  |  | NOS1 | Unknown |
|  |  |  | NOS3 | Activation |
|  |  |  | NOS3 | Unknown |
|  |  |  | ODC1 | Unknown |
|  |  |  | SOD1 | Activation |
|  |  |  | SOD1 | Repression |
| 2 | MYC | v-myc myelocytomatosis viral oncogene homolog (avian) | IL6 | Unknown |
|  |  |  | ODC1 | Activation |
|  |  |  | ODC1 | Unknown |
| 3 | DDIT3 | DNA-damage-inducible transcript 3 | IL6 | Activation |
|  |  |  | NOS3 | Repression |
| 4 | XBP1 | X-box binding protein 1 | IL6 | Unknown |
|  |  |  | NOS2 | Unknown |
| 5 | JUN | jun proto-oncogene | IL6 | Activation |
|  |  |  | IL6 | Unknown |
|  |  |  | NOS2 | Activation |
|  |  |  | NOS2 | Repression |
|  |  |  | NOS2 | Unknown |
|  |  |  | NOS3 | Unknown |
| 6 | AR | androgen receptor | ARG1 | Activation |
|  |  |  | ARG2 | Activation |
| 7 | JUND | jun D proto-oncogene | IL6 | Activation |
|  |  |  | NOS2 | Activation |
| 8 | KLF4 | Kruppel-like factor 4 (gut) | IL6 | Activation |
|  |  |  | ODC1 | Repression |
|  |  |  | SOD1 | Repression |
| 9 | STAT1 | signal transducer and activator of transcription 1, 91kDa | IL6 | Repression |
|  |  |  | NOS2 | Activation |
|  |  |  | NOS2 | Unknown |
| 10 | EGR1 | early growth response 1 | IL6 | Repression |
|  |  |  | SOD1 | Activation |
| 11 | PPARD | peroxisome proliferator-activated receptor delta | CAT | Activation |
|  |  |  | SOD1 | Activation |
| 12 | APC | adenomatous polyposis coli | NOS2 | Activation |
|  |  |  | ODC1 | Repression |
| 13 | NFE2L2 | nuclear factor (erythroid-derived 2)-like 2 | CAT | Activation |
|  |  |  | SOD1 | Activation |
| 14 | XBP1 | X-box binding protein 1 | IL6 | Unknown |
|  |  |  | NOS2 | Unknown |
| 15 | RELA | v-rel reticuloendotheliosis viral oncogene homolog A (avian) | IL6 | Activation |
|  |  |  | IL6 | Repression |
|  |  |  | IL6 | Unknown |
|  |  |  | NOS1 | Activation |
|  |  |  | NOS2 | Activation |
|  |  |  | NOS2 | Unknown |
|  |  |  | NOS3 | Activation |
| 16 | CREB1 | cAMP responsive element binding protein 1 | IL6 | Activation |
|  |  |  | ODC1 | Unknown |

**Figure 1:**
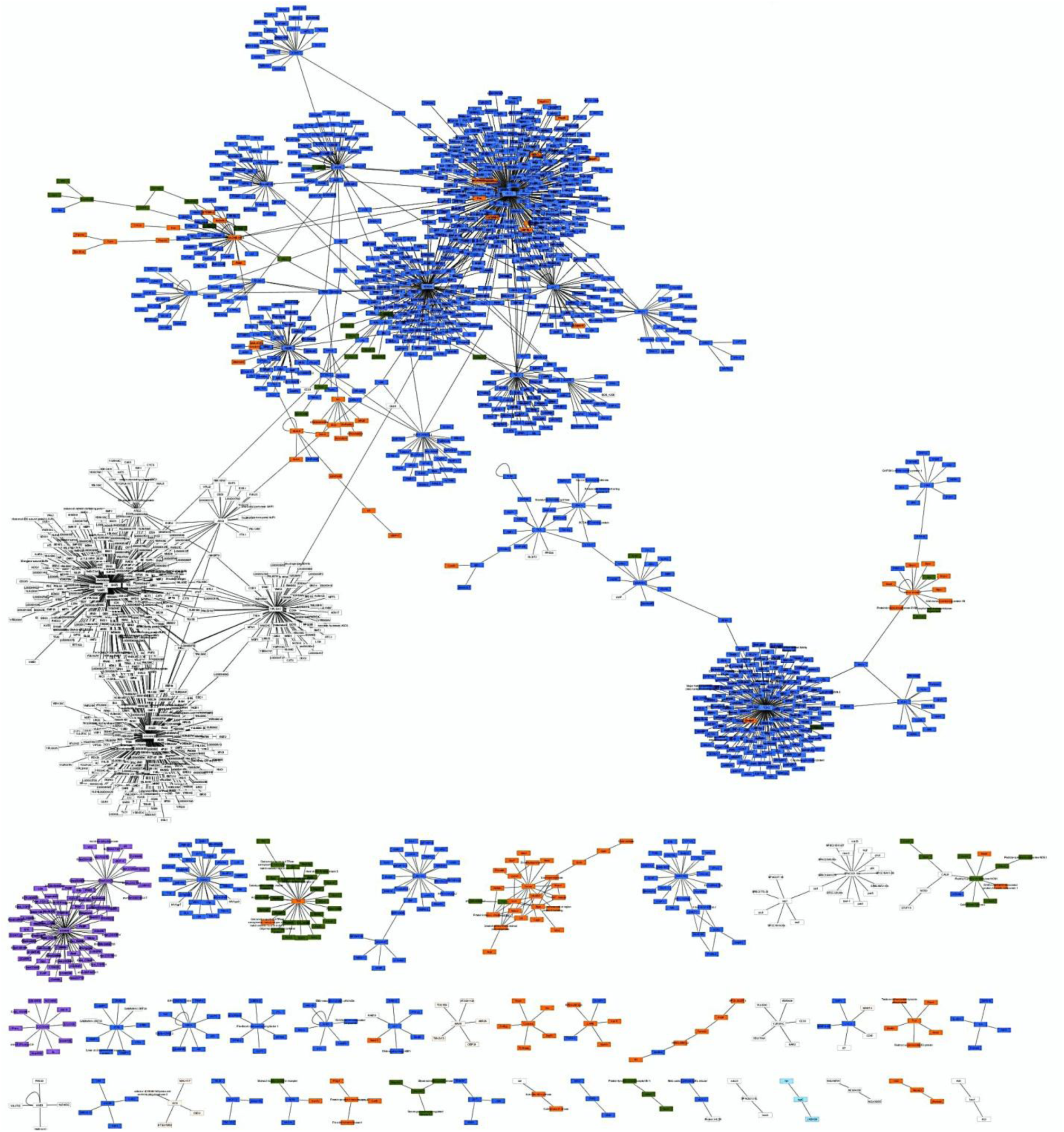
Protein-protein interaction network of nor-NOHA and NOHA associated genes. It contains 1996 nodes (proteins) and 2132 edges (interactions).

**Figure 2:**
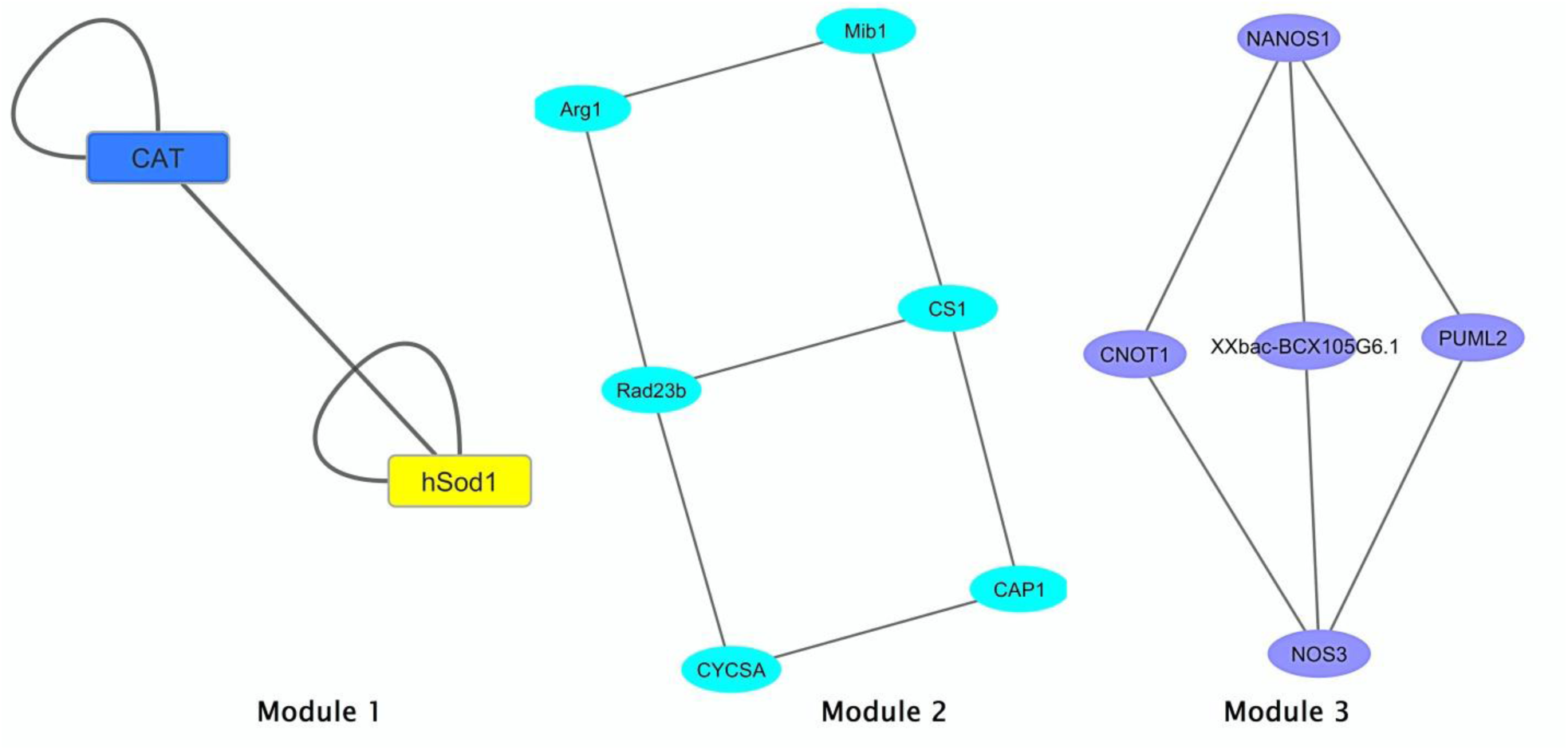
Module 1, 2 and 3 derived using MCODE plug-in in Cytoscape with parameters set as degree cut-off = 2, maximum depth = 100, node score cut-off = 0.2, and -core = 2.

**Figure 3:**
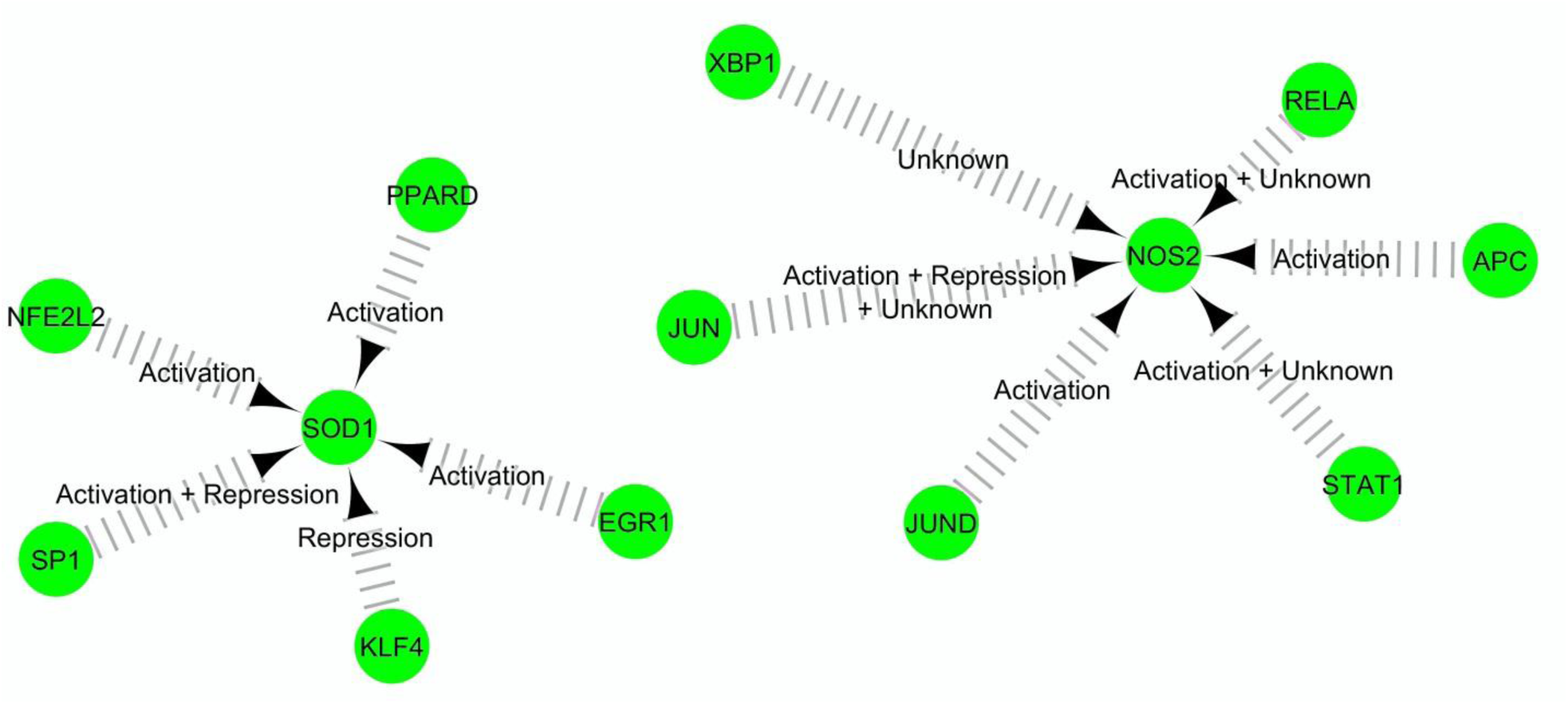
Transcriptional regulatory network of hub proteins SOD1 and NOS2.

Superoxide dismutase has long been considered a target for the selective killing of cancer cells [22]. It has also been labeled as body’s natural cancer fighter [23]. SOD1 however has been only recently been proposed as a novel target for cancer therapy [24] and its role in cancer is also being revealed [25]. More recently being elucidated is the role of AMD1 in cancer which has been found to be upregulated in human prostate cancer [26]. NOS2 is another one which has been recently being seen as a emerging target for cancer treatment [27][28].

## Conclusion

To conclude, a total of 19 nor-NOHA and NOHA-associated genes have been identified. SOD, SOD1, AMD1, and NOS2 have been revealed as key players in nor-NOHA and NOHA mediated pathways. Interestingly, these proteins have been shown to be associated with cancer in other studies as well. Therefore, this study has further validated the arguments for targeting them for treating cancer.

